# Towards a unified resource for transcriptional regulation in *Escherichia coli* K-12: Incorporating high throughput-generated binding data within the classic framework of regulation of initiation of transcription in RegulonDB

**DOI:** 10.1101/219006

**Authors:** Alberto Santos-Zavaleta, Mishael Sánchez-Pérez, Heladia Salgado, David A. Velázquez-Ramírez, Socorro Gama-Castro, Víctor H. Tierrafría, Stephen J.W. Busby, Xin Fang, Bernhard Palsson, James Galagan, Julio Collado-Vides

**Affiliations:** Centro de Ciencias Genómicas, Universidad Nacional Autónoma de México; School of Biosciences, University of Birmingham, Birmingham B15 2TT, United Kingdom; Department of Bioengineering, University of California at San Diego; Center for Biosustainability, The Technical University of Denmark; Department of Biomedical Engineering, Boston University, Massachusetts

## Abstract

Our understanding of the regulation of gene expression has been strongly benefited by the availability of high throughput technologies that enable questioning the whole genome for the binding of specific transcription factors and expression profiles. In the case of genome models, such as *Escherichia coli* K-12, this knowledge needs to be integrated with the legacy of accumulated genetics and molecular biology pre-genomic knowledge in order to attain deeper levels in the understanding of their biology. In spite of the several repositories and curated databases, there is no effort, nor electronic site yet, to comprehensively integrate the available knowledge from all these different sources around the regulation of gene expression of *E. coli* K-12. In this paper, we describe a first effort to expand RegulonDB, the database containing the rich legacy of decades of classic molecular biology experiments supporting what we know about gene regulation and operon organization in *E. coli* K-12, to include the genome-wide data set collections from 25 ChIP and 18 gSELEX publications, respectively, in addition to around 60 expression profiles used in their curation. Three essential features for the integration of this information coming from different methodological approaches are; first, a controlled vocabulary within an ontology for precisely defining growth conditions, second, the criteria to separate elements with enough evidence to consider them involved in gene regulation from isolated sites, and third, an expanded computational model supporting this knowledge. Altogether, this constitutes the basis for adequately gathering and enabling the comparisons and integration strongly needed to manage and access such wealth of knowledge. This version of RegulonBD is a first step toward what should become the unifying access point for current and future knowledge on gene regulation in *E. coli* K-12. Furthermore, this model platform and associated methodologies and criteria, can well be emulated for gathering knowledge on other microbial organisms.

## INTRODUCTION

Equivalent to the role that the elucidation of the structure of DNA had in the foundation of modern genetics, the set of concepts of transcription factor binding sites (TFBSs) affecting the activity of promoters that transcribe transcription units, operons and regulons, constitute the foundation for how we think about gene regulation in microbial organisms, and with some modifications, also in higher organisms. These concepts were the product of research in *Escherichia coli* K-12 during the second half of the last century. It is these concepts that lie behind the computational infrastructure for electronic microbial databases such as RegulonDB, to encode and populate all knowledge that molecular biology has generated from the time of the seminal works by Jacob and Monod until today. More than 20 years of continued curation has meticulously placed, be it a binding site, a transcription factor (TF) and its active conformation, or any other piece of published knowledge on gene regulation, in its corresponding coordinates of the updated complete genome sequence of this bacterium.

However, the emergence of post genomic methodologies has changed the game. We now have whole genome expression profiles for thousands of different conditions (See COLOMBOS and M3D databases, [1, 2] and whole genome identification of binding sites for around 65 TFs, and the numbers continue to increase. As can be seen in Figure 1, we are in the transition of such high-throughput (HT) approaches dominating research as opposed to the more directed molecular biology experiments.

**Figure 1.**
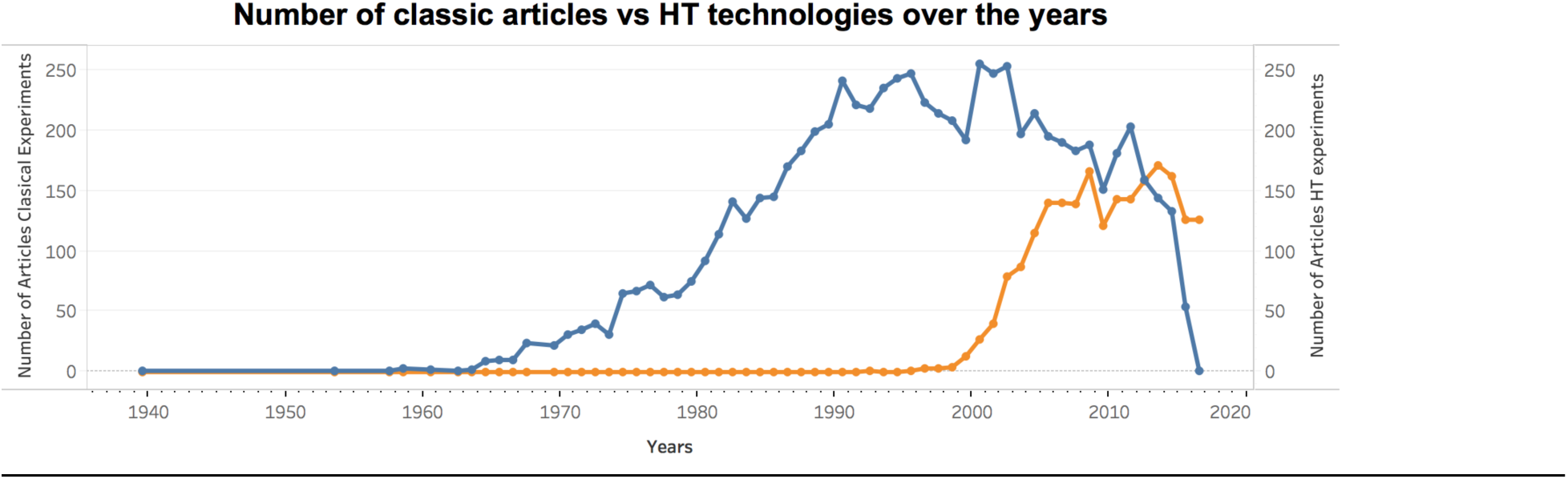
Number of classic articles vs HT technologies over the years of research in *E.coli* K-12. In blue, number of classic articles supporting RegulonDB, and their year of publication versus, in orange, number of HT articles published since 1995 to date, and the years of their publication. HT publications come from searches made in PubMed, GEO and ArrayExpress as well as Scopus. Total number of articles: 7300 with classic methods, plus 2108 with HT-methods.

In the midst of the accelerated pace of data and experimental information generated in the genomic era, databases and other electronic resources are the major instruments to integrate and facilitate access to the tsunami of data otherwise only incompletely apprehended by individual investigators. Table 1 lists the major databases and repositories with information about the biology of *E. coli* K-12. The two up to date manually curated databases are RegulonDB [3] and EcoCyc [4]. Our team is in charge of curating transcriptional regulation for these two databases. On the other hand, COLOMBOS is the only database with microarray data specific for *E. coli*, containing also similar data for a handful of other microorganisms [1]. Otherwise, HT data is found in the general repositories GEO and ArrayExpress.

**Table 1.**

| Source | Type of knowledge | URL | Updated | References |
| --- | --- | --- | --- | --- |
| RegulonDB | Transcriptional regulation, operons, regulons, gensor units | http://regulondb.ccg.unam.mx | Yes | [3] |
| EcoCyc | Regulation, transport and metabolism | https://ecocyc.org | Yes | [4] |
| GenProtEC | Functions encoded by the *Escherichia coli* K-12 genome | http://genprotec.mbl.edu | No | [5] |
| PortEco | Next-generation data *Escherichia coli* | http://porteco.org | No | [6] |
| GenExpDB | Expression Compendia | https://genexpdb.okstate.edu | No | - |
| EcoGene | *E. coli* K-12 genome and proteome information | http://ecogene.org | No | [7] |
| EchoBASE | Information from post-genomic experiments | http://www.ecoli-york.org | No | [8] |
| Colombos | Expression compendia of bacterial organisms | http://colombos.net | Yes | [1] |
| Bacteriome | Integrating physical (protein-protein) and functional interactions | http://www.compsysbio.org/bacteriome/ | - | [9] |
| EcoProDB | Integrates protein information | http://eecoli.kaist.ac.kr/main.html | - | [10] |
| STRING | Protein-protein interaction network | http://string-db.org | Yes | [11] |
| GEO | Genomics high-throughput data repository | https://www.ncbi.nlm.nih.gov/geo/ | Yes | [12] |
| ArrayExpress | Repository of high-throughput functional genomics experiments | https://www.ebi.ac.uk/arrayexpress/ | Yes | [13] |
| M3D | A resource of microbial gene expression data | http://m3d.mssm.edu | No | [2] |

Years ago, there had been efforts in the US to organize HT data for *E. coli* such as the EcoliHub and its subsequent PortEco version, in addition to EcoliWiki, none of them currently actively maintained [6]. Therefore, an investigator interested in gathering all what is currently known about a particular regulatory system in *E. coli* has to spend time searching in these different resources.

Given that HT methodologies precisely enrich our knowledge on gene regulation and gene expression, expanding the current model behind RegulonDB is a natural step to do. However, this is not a simple task, because of several reasons. First, sometimes HT data challenge the Jacob and Monod paradigm, for instance when supporting evidence for a binding site within a gene coding region, or a promoter site within a noncoding region between two convergent ends of genes, where no transcription initiation is supposed to occur. HT methodologies generate large amounts of what sometimes appear as apparently disconnected pieces; for instance, a study reveals around 14,000 candidate TSSs of which more 11,000 occur within coding regions (around 5,500 in the sense strand and around 5,400 antisense) [14]; similarly, a large number of binding sites within coding regions are not any more a surprise in HT-binding experiments. How much of these TSSs or binding sites are either spurious or functional and participate in roles not directly related to gene regulation is still an open question.

As a consequence, we need a mixed model with room both for the classic complete stories where objects and their interactions make sense, as well as with room for plausible disconnected objects. First, the data should be available both in a structured way when possible, but also in a non-prejudiced agnostic way so the user can make his own decisions. Second, we need to implement tools and criteria to identify experiments performed under similar conditions. An ontology and its corresponding controlled vocabulary for precisely defining growth conditions is part of our efforts in this direction, as described in (Tierrafria et al., submitted). This is the basis for merging our classic curation with the one presented here of HT-binding experiments, together with expression profiles to identify the effect of binding to construct a regulatory interaction. Third, we need to define additional evidence codes for different types of HT experiments, together with the limits that define when there is sufficient information for a new regulatory interaction, or any other piece of evidence that contributes plausible regulatory processes, as opposed to scattered elements with not enough support for their interpretation as functional elements of gene regulation. Finally, we had to define what features and how to display HT generated binding sites and regulatory interactions, consistently with those already existing. Altogether, this constitutes the basis for adequately gathering and enabling the comparisons and integration strongly needed to manage current knowledge about transcriptional regulation in *E. coli*. We present here the first version of a more complete integration of HT-binding experiments (ChIP and gSELEX) with the previously curated literature.

## MATERIALS AND METHODS

Figure 1 was generated by identifying the date of publication for all 7356 papers that had been curated in RegulonDB. The HT publications were obtained by searching in PubMed, GEO and ArrayExpress as well as Scopus, from year 1995 to 2017. A total of 2108 HT-papers were obtained. Of these, around 50 papers were processed in order to extract all peak sequences or regions identified by the HT-binding methods. Frequently, these papers include additional experimental characterization for a subset of the sites with EMSA, footprinting analysis, and used bioinformatics tools, mostly using position weight matrices (PWMs) for the transcription factor binding sites (TFBSs) to precisely identify the binding sites in the sequences of the peak regions. Curation of this literature extracted from each publication included the following metadata: the strain, the growth condition, number of targets, name of the TF, methodology used (ChIP-X or gSELEX) or RNA-seq, and its evidence code, additional techniques used to further identify the binding sites; as well as links to the files, when available in the repositories of GEO or ArrayExpress. As mentioned, the growth condition and strain are described using the controlled vocabulary defined in (Tierrafria et. al.,submitted). As explained in the section on the curation of HT literature, the products of curation are added in RegulonDB either together with the classic curation, or as separate datasets. For those added to the classic curation the information includes the sequence of the TFBS, its evidence and reference, its precise coordinates; the peak coordinates start-end, with its evidence; the effect or function (activation, repression, or dual) with its evidence and method, we also indicate if the effect was identified by the authors or by us; and the regulated gene. Information on the peak sequences is contained in the datasets. Remember that once the DNA sequences brought by the antibody are sequenced, these are then mapped to the genome sequence, defining the sequence peaks or regions, in these experiments usually on the range of 500 to 200 nucleotides. We refer to them as peak sequences. A subsequent step is that of identifying the potential precise binding site for the given TF. Most of the time this is currently done by running alternative bioinformatics methods that use known position weight matrices (PWMs) within those regions, such as MEME [15], dyad analysis or other similar methods [16], although alternative methods exist [17, 18]. We gathered the method used, as well the evidence according to the notation used in RegulonDB that expands the one from the Gene Ontology consortium (http://regulondb.ccg.unam.mx/evidenceclassification).

In several cases the sequences that result from the peak calling algorithms were provided without identification of the precise binding site. In those cases, the curator team used the PWM available in RegulonDB for the given TF (see: http://regulondb.ccg.unam.mx/external_data/MatrixAlignment/results/) to search in the peak sequences using the threshold parameters adequate for each TF. For all identified TFBSs we searched for evidence of change in expression of the downstream gene and TU in COLOMBOS and available RNA-seq experiments. A minimal change of 2-fold and a p-value of 0.05 was considered to indicate change of expression.

## RESULTS

This paper focuses in the literature of HT binding experiments, and currently we have limited our curation to incorporate those objects that are processed and proposed in the publications by the authors. To do this, we implemented a simple strategy that separates objects (sites, promoters, interactions) that satisfies a set of criteria (of confidence and interpretability, see below), and were uploaded in RegulonDB together will all existing knowledge. When these criteria are not satisfied, then we simply offer the data as datasets (see: http://regulondb.ccg.unam.mx/central_panel_menu/downloads_menu.jsp). The major difference is that datasets are not equally browsable or displayed within RegulonDB as explained below.

### Curation of HT literature in RegulonDB

As reported in our publications on progress in RegulonDB, we had curated some few papers from HT experiments years ago. The first dataset that has been available are transcription start sites (TSSs) identified by Illumina sequencing of 5´triphosphate enriched transcripts by the group of Morett [19]; later in 2015 we initiated the curation of binding sites from gSELEX (CRP, H-NS and LeuO), and from ChIP-exo (GadE, GadW, GadY, OxyR and SoxS), as well as the dataset of TSSs by the group of Gisela Storz [14]. We are now including curated sites and made a separate section so that the user can quickly see either the set coming from HT experiments, together and/or separated from those coming from classic methods. Furthermore, we have initiated important modifications to the computational model of RegulonDB, together with a controlled vocabulary for growth conditions, which all together prepare us for a constant and eventual up to date curation of all this literature and content. We extracted publicly available binding sites for 42 different TFs from experiments performed in *E. coli* K-12 by ChIP (ChIP-chip, ChIP-seq y ChIP-exo), or genomic SELEX (gSELEX) by the group of Ishihama performed in the *E. coli* str. K-12 substr. W3110 [20], (indicated as a note in RegulonDB); as well as RNA seq and microarrays information contained in those papers. Curation of this literature included extracting the metadata (see methods) that contain all relevant information of the biology (TF and growth conditions) as well as links to the data if it is found in standard repositories, and relevant information as detailed in the methods section. A total of 18 papers with gSELEX data were curated, 14 from ChIP-chip, 5 from ChIP-seq and 4 with ChIP-exo. The summary of all curated knowledge from HT methodologies currently available in RegulonDB is shown in Table 2. This is an important first step, additional data is being continually curated in order to reach an up to date level equal with the classic literature.

**Table 2.** Summary of all curated knowledge from HT methodologies currently available in RegulonDB.

| Methodologies | Number of articles | Number of TFs | Name of the TFs |
| --- | --- | --- | --- |
| gSELEX | 2 previous work | 3 | CRP, H-NS and LeuO |
|  | 18 this work | 18 | AscG, BasR, CitB, Cra, CsgD, Dan, DpiA, LeuO, Lrp, NemR, OmpR, PdhR, PgrR, RcdA, RstA, RutR, SdiA, and SutR |
| ChIP-chip | 1 previous work | 1 | PurR |
|  | 14 this work | 14 | ArcA, ArgR, CRP, Fis, FNR, H-NS, IHF, LexA, Lrp, NsrR, RpoD (Sigma70), RpoH (Sigma32), RutR, and TrpR |
| ChIP-exo | 2 previous work | 6 | GadE, GadW, GadX, OxyR, SoxS and SoxR |
|  | 6 this work | 4 | ArgR, Fur, OmpR, and UvrY |
| ChIP-seq | 5 this work | 8 | CsiR, FNR, Fur, H-NS, Nac, OmpR, RpoD (Sigma70), and RpoS (Sigma38) |
| Methodologies | Number of articles | Number of TSSs | Dataset in RegulonDB |
| TSS determination | 2 previous work | 5,197 | http://regulondb.ccg.unam.mx/menu/download/high_throughput_datasets/index.jsp |
|  |  | 1,806 | http://regulondb.ccg.unam.mx/menu/download/high_throughput_datasets/index.jsp |
|  | 1 previous work | 14,000 | http://regulondb.ccg.unam.mx/menu/download/high_throughput_datasets/index.jsp |

### Criteria to combine classic and HT supported data

In the role of editors of the knowledge on gene regulation in *E. coli*, the best decision we can make is to offer users the best possible integration of data and information, clearly indicating their experimental method and reference. The challenge of the classic paradigm of gene regulation with the scattered data from HT experiments is solved in practice by separating two sets as the product of our curation: those pieces of knowledge (TFBSs) with enough additional evidence to support their functional role in gene regulation are added to the bulk of existing knowledge, whereas all binding sites for which not enough information is found yet to know if the bound TF has a role in gene regulation, are kept in separate datasets. Additionally, experiments kept in datasets are those that support a given DNA region in the genome that is usually much larger than TFBSs, such as peak regions, or regions from SELEX experiments, but which do not have a precise TFBS identified.

Users can download and combine both the information available within the classic model of RegulonDB with any of the available datasets, and we plan on implementing additional tools in the future that will facilitate their comparison, visualization and processing. As we implement them in the future, the decision of what is being added to the core of knowledge and what is left as datasets will be in practice less relevant.

It should be clear that in order to find the additional evidence we first curated what the authors did to support their binding sites as elements that are playing a role in gene regulation, frequently by performing additional experiments. And in a more active curation, we also searched in other publications and datasets, for evidence needed to suggest the effect on regulation, activating or repressing transcription. We specifically combined data from gene expression generated by RNA-seq and/or microarray experiments, with data from TF DNA binding experiments. To do so, we used our parallel work of mapping growth conditions in RegulonDB with growth conditions in COLOMBOS. Such a mapping and definition of a controlled vocabulary is an enormous task not yet completed, but in our coordinated work, we made sure the conditions present in our meta-curation for HT experiments were included. For details see (Tierrafría et al., submitted).

The central question then is: What is the minimal evidence that supports a site found by either any ChIP type of experiment (ChIP-seq, ChIP-exo, ChIP-chip), or by gSELEX to have a functional role in gene regulation? First of all, as mentioned, we consider that the binding site sequence has to be identified, otherwise the TF-target gene could be indirect. The set of cases that we considered that support a regulatory interaction in order of supported evidence, are the following. The more solid ones are those with a sequence identified for binding of a TF, frequently identified by a computational search in the peak sequence, and the effect on regulation suggested by an observed change in gene expression. We assigned the effect (activator, repressor or dual effect) determined for the regulated gene or transcription unit. If the regulatory interaction and TFBS did not exist in RegulonDB, it was added as a new site and a new regulatory interaction. If it already exists, then the new evidence was added to the existing regulatory interaction.

In case the authors did not identify the precise TFBS, we used the PWMs in RegulonDB and searched for a binding site in the sequence, and only when found, it was added as a regulatory interaction. The following cases are considered to have insufficient information to know if they play a role in gene regulation. Those where the binding site identified but in the absence of any evidence to assign an effect and a regulated gene. In some cases, the corresponding expression experiment has been performed, but there is no evidence of change in expression of the downstream gene. The possible reasons for this could be: Inactive conformation of the TF or co-regulation missing in the condition studied, or effectively the protein binds but has no role in transcriptional regulation. For the time being, we have decided that peak sequences with or without a binding site identified, that fall in regions of the genome where no transcription is expected, such as within a coding region, or within a convergent region surrounded by the end of two genes, were not further analyzed, and can be accessed only as datasets. We are aware that additional work can be done, for instance by searching for transcription start sites nearby, curating antisense transcription (currently in datasets), by systematically searching for plausible expression in RNA seq experiments, and curating cases of TFBSs within genes with a regulatory effect (See the site for Nac inside the GadE gene, and Tables 1 and 2 in Aquino et al., 2017) [21].

In addition to the evidence code and the method, we show our classification of evidence as either confirmed, strong or weak. Evidence codes comes from the Gene Ontology consortium shared in our curation both by RegulonDB and EcoCyc. In order to facilitate the processing by the user of the diversity of evidence codes, in RegulonDB we classify them in three classes: “confirmed” when they have more than one independent solid evidence, “strong” for cases supported by physical evidence, and “weak” otherwise (such as a computational prediction). Objects with multiple independent weak evidence are upgraded to strong; a detailed explanation is found in http://regulondb.ccg.unam.mx/evidenceclassification which is the product of [22]. Note that we always include the precise evidence codes for more detail, also in case users do not like this classification of types of evidence unique to RegulonDB. A summary of the results of this curation is shown in Table 3. We will call HT-supported regulatory interactions those sites that satisfy the minimal criteria outlined, and “HT-binding sites” those that were left as datasets.

**Table 3.** Summary of curated HT-regulatory interactions and sites in datasets

| Complete data for upload to RegulonDB |  |  |  |
| --- | --- | --- | --- |
| TF | Number of interactions | PMID | HT methodology |
| ArgR | 85 | 25735747 | ChIP-exo |
| ArcA | 135 | 24699140 | ChIP-chip |
| FNR | 38 | 24699140 | ChIP-chip |
| Lrp | 76 | 19052235 | ChIP-chip |
| Fur | 57 | 25222563 | ChIP-exo |
| OmpR | 19 | 28526842 | ChIP-exo |
| As datasets |  |  |  |
| TF | Number of interactions | PMID | HT methodology |
| ArgR | 426 | 22082910 | ChIP-chip |
| ArcA | 143 | 24699140 | ChIP-chip |
| ArgR | 38 | 25735747 | ChIP-exo |
| CRP | 39 | 16301522 | ChIP-chip |
| CsiR | 126 | 28061857 | ChIP-seq |
| Fis | 228 | 16963779 | ChIP-chip |
| FNR | 137 | 17164287 | ChIP-chip |
| FNR | 227 | 23818864 | ChIP-seq |
| FNR | 186 | 24699141 | ChIP-chip |
| Fur | 473 | 26670385 | ChIP-seq |
| Fur | 87 | 25222563 | ChIP-exo |
| H-NS | 101 | 16963779 | ChIP-chip |
| H-NS | 53 | 21097887 | ChIP-seq |
| IHF | 155 | 16963779 | ChIP-chip |
| LexA | 69 | 16264194 | ChIP-chip |
| Lrp | 67 | 19052235 | ChIP-chip |
| Nac | 537 | 28061857 | ChIP-seq |
| NsrR | 83 | 19656291 | ChIP-chip |
| OmpR | 68 | 28061857 | ChIP-seq |
| OmpR | 22 | 28526842 | ChIP-exo |
| RpoD (Sigma70) | 1214 | 16109958 | ChIP-chip |
| RpoD (Sigma70) | 528 | 16301522 | ChIP-chip |
| RpoD (Sigma70) | 199 | 19706412 | ChIP-chip |
| RpoD (Sigma70) | 691 | 23818864 | ChIP-seq |
| RpoH (Sigma32) | 82 | 16892065 | ChIP-chip |
| RpoH (Sigma32) | 44 | 20602746 | ChIP-chip |
| RpoS (Sigma38) | 78 | 26020590 | ChIP-seq |
| RutR | 20 | 18515344 | ChIP-chip |
| TrpR | 17 | 22082910 | ChIP-chip |
| UvrY | 340 | 26673755 | ChIP-exo |

### Display in RegulonDB

All these curated HT-supported regulatory interactions are now present within RegulonDB, and can be found in the regulon page of the corresponding TF. The most direct way to access them is to type the TF name followed by “regulon”, go to the link of the regulon, and display the TF regulon page. In that page, there is a table with all TFBSs, now including those that come from HT experiments. Table 3 describes all TFs with HT supported binding sites in the current version of the database. Furthermore, via the Downloads main page menu, high-throughput datasets can be selected and then select any of the TF specific HT-binding datasets. Both of them (individual HT-supported TFBSs, and specific datasets) can be browsed by searching on growth conditions for instance, particularly using their contrasting experimental vs control condition change. Additionally, as already mentioned, a search using the controlled vocabulary for growth conditions will show both the structured data as well as the link to the datasets. We are working to display any dataset as a track in our browser, which will enable the direct comparison with, for instance, information coming from classic experiments, and with any other annotations available in RegulonDB.

## DISCUSSION

As mentioned before, we do not want to dilute the predominantly highly confident knowledge that comes from classic experimental methods aimed at identifying individual objects or interactions, with the massive but more fragmented knowledge that HT methodologies produce [3], which by their proper nature involve several layers of experimental treatments and subsequent processing by bioinformatics and statistical methods. Thus, not only the experimental methodology varies, but also the bioinformatics programs, and the selection of thresholds used in the different processing steps. Nonetheless, as shown in Figure 1, the tendency of the literature is the continuous and more dominant use of HT-methods in research, proving the urgent need for this expansion of RegulonDB. This requires to modify several components of our system, starting by a computational model with much more precise encoding of the distinct, almost elementary components that constitute the knowledge of gene regulation: We now require evidence, methods and reference for the binding site of a given TF, and separately for its effect on a regulated gene or promoter; we need to indicate the expression profile experiment that supported a change in expression of the (candidate) regulated gene; we also distinguish which piece comes from the literature and which one comes from our own active curation. It is important to note that even classic experiments generate, by the proper nature of experimental work, pieces of evidence that are gradually constructed to generate the picture necessary to make sense of it. For instance, the gene regulated by a TF is frequently identified by transcriptional constructions with a reporter gene. Strictly speaking this evidence only supports the fact that RNA polymerase proceeds into transcription downstream of the promoter; whether it transcribes *in vivo* only the first downstream gene or the complete transcription unit requires identification of such transcript under precisely the same control and regulated conditions.

Our controlled vocabulary and collection of features generically called “growth conditions” also contribute to higher precision by annotating the strain or genetic background used in the experiment, as well as growth conditions minimally required for their replicability. We believe that as we advance in this decomposition down to the “elementary pieces of knowledge” as they come from experiments, we will be better prepared to incorporate experiments from the new methodologies that will continue to emerge in the future. This expanded model affects both the internal structure, the tools for curation and the display for users to access the data. In this paper, we focused essentially on HT alternatives that identify binding sites for transcriptional regulators at a genomic level. These experiments identify the bound sites in the genome, some of which may have a role *in vivo* affecting gene regulation, but others may have no role at all in affecting transcription, and therefore even their name of “transcription factor binding sites” may be misleading.

**Figure 2.**
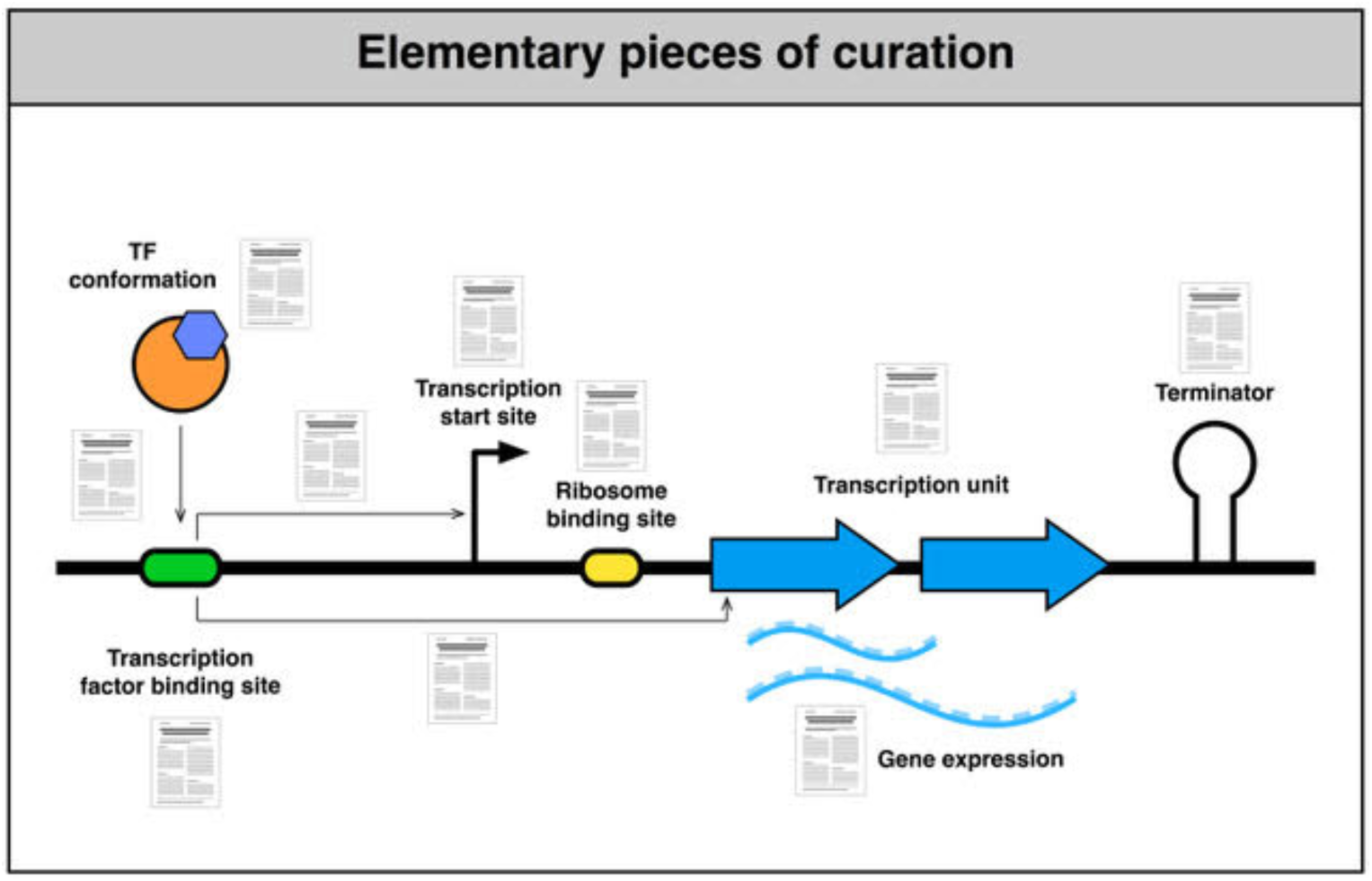
Elementary pieces of curation. As new methods emerge we need to separately curate evidence and references for each elementary piece of knowledge that together support our understanding. We have here separated evidence for the binding of TFs, evidence for the effect on transcription either on a known promoter, or on a target gene or TU without knowing the promoter.

The strategy used both in the computational model and the display of knowledge enables users to decide if they want to see either the knowledge that comes from molecular biology experiments, that from HT-methods, or both.

We consider the work here presented as a first version of what we envision is a long-term project required to include the many components required. Certainly, there is wide room for improvements. Many more analyses can be implemented in cross-comparisons of the increasing HT-datasets, so that new correlations may emerge. In this sense, the curation presented here has only used the assignment of effect of the TFBS by searching in the biologically adequate expression profile to see if the change of expression of the downstream gene was observed, where by “adequate profile” we mean the comparable growth condition and strain. But in fact, many more analyses can be performed. For instance, it will be, it will be useful to offer datasets with partial knowledge that regulate gene expression by unknown mechanisms, such as those occurring within coding regions [21]. Additional programs need to be implemented to search for all binding sites, if there are TSSs found nearby, including the thousands present in our datasets. The relative distance of TFBSs to its regulated TSS is known to correlate with the activating or repressing function [23, 24]; some sigma factors are associated to particular conditions, like stress or heat shock. All of this and more, provides food for pipelines to be implemented for a more automatic and periodic update in the generation of evidence for gene regulation. This suggests a new type of “bioinformatics biocuration” with a more active work gathering evidence across multiple publications, experiments and evidence to gradually reconstruct the different elements and interactions required in our understanding of the regulation of transcription initiation and to distinguish those involved in gene regulation by unknown mechanisms, as well as those that may have different roles associated to their binding in yet unknown processes in evolution.

## ACKNOWLEDGEMENTS

We acknowledge technical support by Víctor Del Moral, Kevin Alquicira Hernández, Jair Garcia Sotelo, and Alfredo Hernández. This work was supported by UNAM, by the National Institutes of Health NIGMS [grant number R01GM110597-3]; and by FOINS CONACyT Fronteras de la Ciencia [project 15].

